# Parasite gene flow in riverine habitats: ascertaining the roles of stream drift, river bifurcations and host dispersal

**DOI:** 10.1101/2022.09.21.508869

**Authors:** Mary J. Janecka

## Abstract

Determining the factors that shape parasite gene flow across complex landscapes is central to understanding the coevolutionary process. In rivers, unidirectional currents, stream drift, may facilitate downstream parasite dispersal, while bifurcating branches may cause population subdivision among branches. The generative habitat processes in rivers can potentially interact with host dispersal to determine gene flow within the aquatic ecosystem. We examined the population genetic structure and gene flow of a trematode infecting semi-aquatic snakes to determine the relative contributions of stream drift, river bifurcations and host dispersal in shaping parasite gene flow in three connected riverine ecosystems. We found the strongest population structure immediately below a recently constructed reservoir at the confluence of the two rivers, with mild structure between one out the the three reaches of the river. Patterns of isolation by distance along linear pathways were not uniform, despite similar path network path lengths. We found the strongest evidence for isolation by distance associated with the river bifurcation. The comparison of terrestrial versus within river network dispersal indicates that parasite transmission between branches occurs along river networks. Short-distance terrestrial dispersal however may be important along some linear networks. Our results highlight the complexity of host-habitat interactions shaping parasite gene flow and the need for empirical data from natural systems to develop accurate models of parasite transmission in rivers.

## 1. Introdution

Estimating gene flow is a key population parameter essential to understanding local adaptation, population connectivity and the evolutionary trajectory of a species (Gandon and Michalakis 2002; Gandon et al. 1996; McCoy et al. 2003; Johnson et al. 2020). Realized gene flow across the landscape will be the results of complex interactions between life history, size and dispersal ability of the organism and physical features of the landscape that act as barriers to dispersal (Crispo et al. 2006; Criscione and Blouin 2004; Pilger et al. 2017)

An important difference between free-living and parasitic organisms is that parasites are subdivided among hosts, and thus parasite gene flow may be affected by host dispersal in addition to land-scape scale processes (Gorton et al. 2012). As host-parasite co-evolutionary dynamics are dependent on the relative migration rates between host and parasite, geographic variation in dispersal of infected hosts, and thus the parasites themselves, can also interact with complex landscapes to fundamentally influence host-parasite evolutionary, demographic and ecological trajectories (Fernandes et al. 2019; Gomulkiewicz et al. 2000; Johnson et al. 2020) Thus, understanding the factors that shape parasite gene flow across complex landscapes is a central component in understanding the coevolutionary process. With their small size and short-lived free-living stage, dispersal for parasites with complex life cycles, the most mobile host in the life cycle is predicted to exert the most influence on parasite dispersal (Prugnolle *et al*. 2005; Louhi *et al*. 2010; Blasco-Costa *et al*. 2012). When applied to intraspecific genetic variation, the autogenic-allogeneic hypothesis predicts that autogenic parasites, who complete their life cycle entirely in aquatic ecosystems, will have lower gene flow among subpopulations and by extension, greater population structure than autogenic parasites, whose hosts can also disperse across terrestrially or aerially (Esch et al. 1998; Criscione and Blouin 2004). Previous research has found support for this hypothesis in comparisons between fish and avian or mammalian definitive hosts (Criscione and Blouin 2004; Blasco-Costa and Poulin 2013), however, this dichotomy fails to capture the nuance and complexity of aquatic and semi-aquatic freshwater organisms who very both in dispersal distance and potential for terrestrial dispersal.

River systems are unique landscape configurations characterized by their dendritic structure and unidirectional stream flow that have the potential to shape spatial patterns of dispersal and gene flow for the organisms constrained within them (Blanchet *et al*. 2020). Altitudinal gradients generate asymmetric, downstream water flow, while the repeated arborescent bifurcations create a spatial hierarchy of unique landscape pathways along branches and nodes. The unidirectional stream drift of river systems mediates the down-stream biased asymmetric migration rates for fluvial organisms, potentially generating a pattern of downstream increases in genetic diversity, and isolation by distance along linear pathways of the river when gene flow is similarly unidirectional (Blanchet *et al*. 2020). Stream drift has been documented to be a major force behind the dispersal of invertebrate organisms including snails, as well as small fish and tadpoles that serve as intermediate hosts for many trematodes, resulting in many species exhibiting the characteristic down-stream increase in genetic diversity (hereafter DIGD) (Poff *et al*. 1997; Crispo *et al*. 2006; Blondel *et al*. 2019, 2020).

The spatial hierarchy of the branching networks can be characterized as distal or confluent pathways, which may differentially facilitate decreased or increased dispersal corridors (Chiu *et al*. 2020). Dispersal asymmetry can also vary spatially within the riverine network, facilitating gene flow along mainstem corridors while distal headwaters may experience less connectivity and subsequent population subdivision and isolation by distance among metapopulations (Alp *et al*. 2012; Paz-Vinas & Blanchet 2015; Thomaz *et al*. 2016). Theoretical modeling that addresses network connectivity predicts clustered branches and shorter branch lengths will increase connectivity, while increasing branch complexity and long network pathways generate population subdivision (Labonne *et al*. 2008; Alp *et al*. 2012; Paz-Vinas & Blanchet 2015; Paz-Vinas *et al*. 2015; Schmidt & Schaefer 2018; Alther et al. 2021). Short-distance dispersers may exhibit extreme isolation and endisms constrained within branches, isolation by distance in river networks, and downstream-biased migration (Thornton 2007; Campbell Grant *et al*. 2007; Whelan *et al*. 2019). Long-distance dispersal of fully aquatic organisms may still be constrained by river network connectivity between drainages. Other freshwater vertebrates, such as amphibians and reptiles, however, may be capable of out of limited out-of-network movement. The potential for out of network movement of semi-aquatic species may vary, depending on the resilience of the organisms to desiccation and the degree to which the surrounding landscape provides temporary refugia from desiccation (Miller et al. 2015; Ciofi et al 2017; Roe et al. 2003).

Parasites are an integral and ubiquitous facet of river ecosystems. Infection from parasites with riverine or partially riverine life-cycles exact high costs to free-living hosts and can impose alterations to host reproduction, behavior and survival (Anderson & May 1979; Marcogliese & Pietrock 2011; Walsman *et al*. 2021). Riverine parasites also cause significant morbidity and mortality to fish of conservation (Lymbery *et al*. 2020) and commercial concern (Krkosek *et al*. 2007). Unfortunately, our understanding of how river architecture and hosts may interact to influence parasite genetic diversity and transmission in riverine systems is largely unknown (Blasco-Costa *et al*. 2012, 2013; Blasco-Costa & Poulin 2013; Janecka *et al*. 2021b).

The objective of our study was to examine interactions between stream drift, DEN bifurcations, and host dispersal on the distribution of parasite genetic diversity and population subdivision within a simple, bifurcating river system with similar network pathways among branches, i.e., the Colorado and Concho Rivers of west central Texas (Figure 1). The generative river processes of stream drift and DEN’s in shaping parasite genetic diversity and population structure will ultimately depend on complex interactions between the habitat structure, parasite life history, and host dispersal capability. Thus, the stream drift and DEN predicted population genetic patterns for fully aquatic organisms may not apply to riverine parasites that use hosts with high dispersal abilities or semiaquatic hosts that can circumvent barriers experienced by fully aquatic species.

**FIGURE 1.**
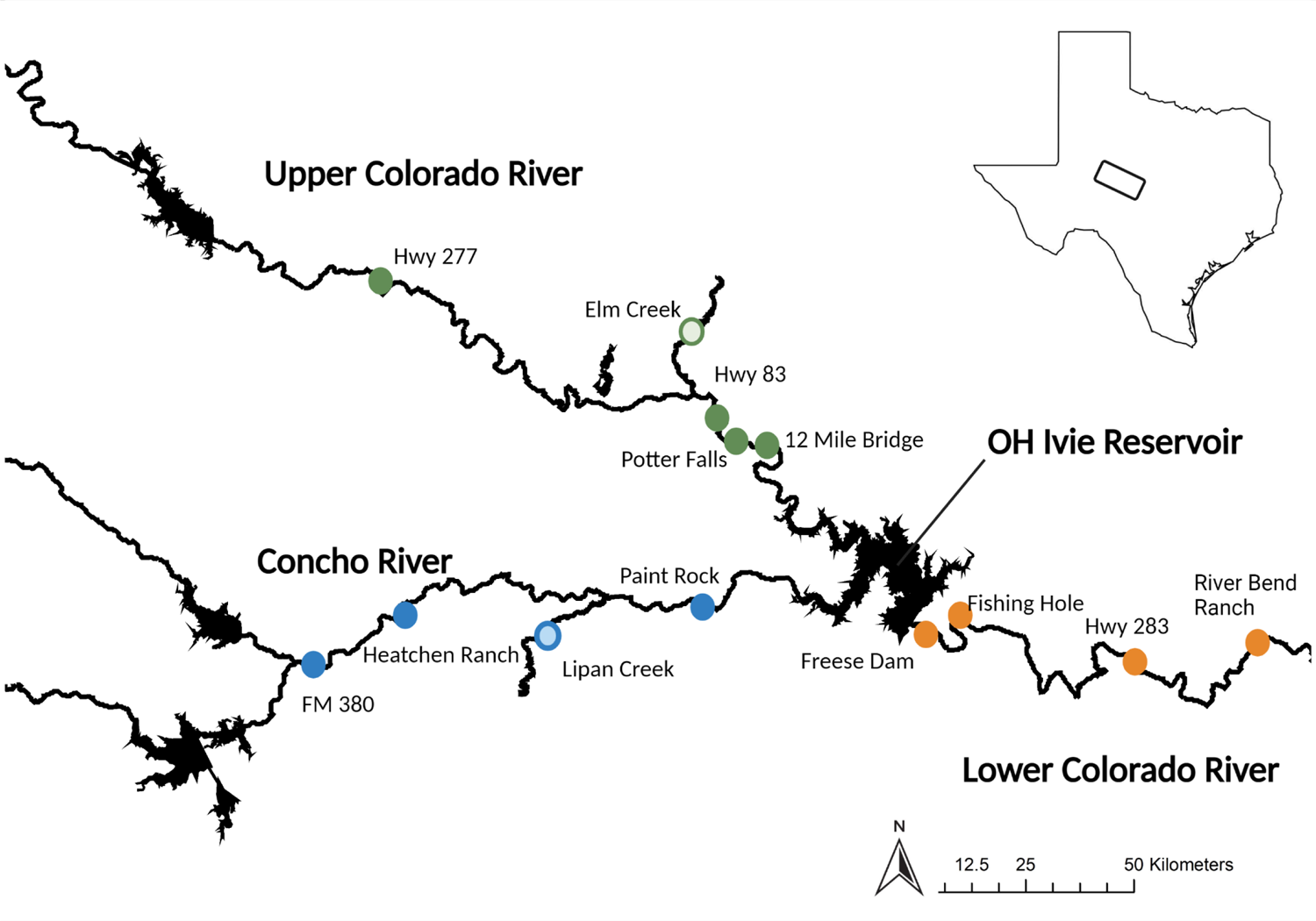
A. Map of sampling sites on the Colorado and Concho Rivers in west central Texas, USA. The Upper Colorado and Concho Rivers form a confluence within OH Ivie Reservoir

Our study parasite was *Renifer aniarum*, a digenean trematode that as sexually mature adults infect the mouths of semi-aquatic water snakes (Figure 2). Snakes become infected through trophic transmission when they consume tadpoles infected with metacercariae (Byrd 1935). Figure 2 shows the cycle for which we note that all the aquatic stages (free-living and parasitic) are likely subject to stream drift and DEN effects as shown in other aquatic invertebrates (Talbot, 1934; Byrd, 1935; Byrd and Denton, 1938; Alp et al. 2012, Whelan et al, 2019). However, the snake final hosts may negate such impacts. *Renifer aniarum* infects the entire water snake community of the Colorado and Concho Rivers: *Nerodia erythrogaster*, *N. rhombifer* and *N. paucimaculata* (Janecka et al. *in prep*). The population structure for two of the snake species has been examined: Janecka et al. (2021) found isolation by distance and population subdivision in *N. paucimatuala*, while Rodriguez et al. (2012) found no population subdivision for *N. erythrogaste*r among the three reaches of the river. The parasite’s use of multiple snake species, especially where one shows no genetic structure among the sampled region, may facilitate its dispersal up stream and across branches of the river and thereby, eliminate or reduce the influence of stream drift and DEN on its population structure.

**FIGURE 2.**
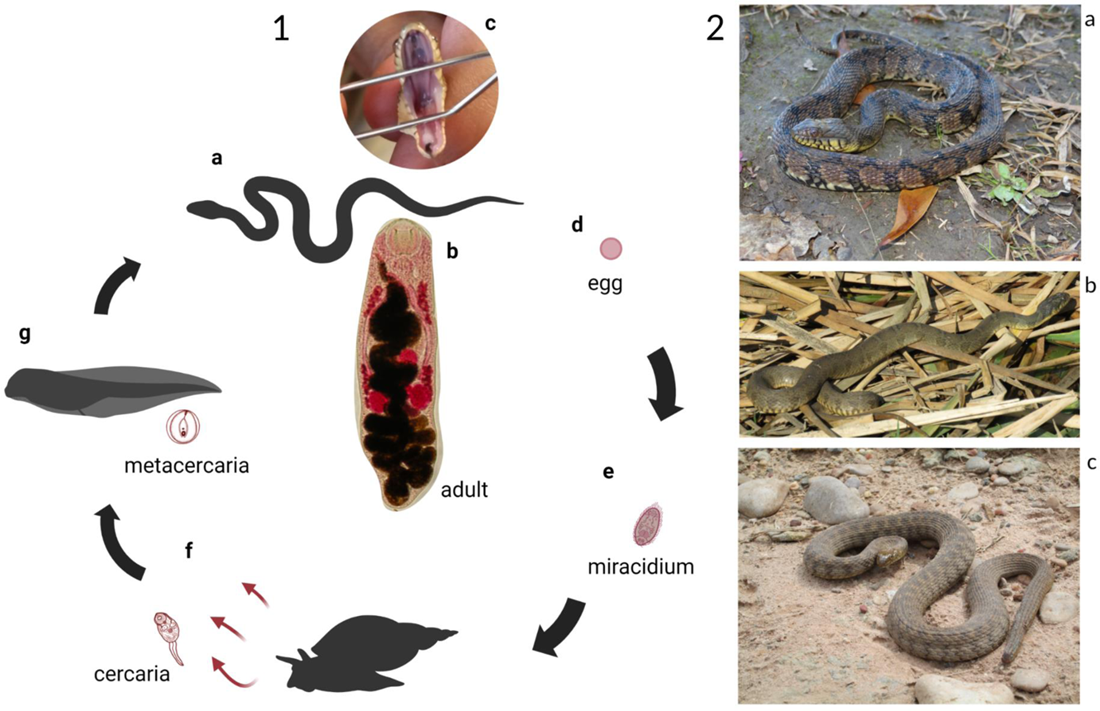
1. Life cycle of *R. aniarum.* Adult flukes infect the mouths of three species of water snake in the study area (a-c). Eggs are passed through the feces (d) and free-living miracidia (e) infect snails (Physa spp) as the first intermediate host. Cercaria exit the snail (f) and encyst on the tails of tadpoles (*Rana* spp*., Hyla* spp., *Psuedacris* spp., *Lithobates* app. and *Anaxyrus* spp.). Snakes become infected when they consume tadpoles with the encysted *R. aniarum* metacercaria. 1.2 Three species of water snake were found to be infected by *R. aniarum* in the study site.

We address three questions to examine the relative influence of riverine habitats and hosts within this system. If the physical characteristics of the river are the primary influence on *R. aniarum* population structure than we predict that one or both of two non-mutually exclusive results would be observed: 1). Is stream drift shaping *R. aniarum* genetic diversity? If yes, then populations will increase in genetic diversity with an increasing distance downstream and exhibit moderate to high genetic structure among subpopulations. Additionally, populations downstream of the confluence of the Concho and Colorado Rivers should be more genetically diverse than upstream populations due to the contributions of the two separate upstream drainages. 2). Does the DEN shape *R. aniarum* population structure? If yes, population subdivision will be associated with the structural features of the river system, specifically among the three reaches of the Colorado and Concho Rivers and the Elm Creek and Lipan Creek tributaries. We also predict isolation by river distance between river branches if parasite dispersal is constrained by river branches. And 3). Do hosts circumvent the barriers to aquatic systems (dams and network branches) via terrestrial dispersal? If yes, then we predict isolation by straight-line distance or no population structure.

## 2. Material and Methods

### 2.1 River system, surveys and sample collection

The Colorado and Concho Rivers are the primary water source for several municipalities as well as irrigation for cattle and livestock in semi-arid west Texas. The Concho and Colorado Rivers form a confluence within O.H. Ivie Reservoir, separating the river system into three reaches, the Upper Colorado River (hear after UCR) above O.H. Ivie reservoir, the Lower Colorado River downstream of O.H. Ivie reservoir, and the Concho Rivers (CHR), forming a relatively simple bifurcation with similar river distances (∼ 175 km) among the three distal sites of the bifurcation. Eleven sites were selected to encompass locations on the three reaches of the river and two tributaries (13 total) (Figure 1). Site location was based in part on accessibility, network placement and to coincide with post-delisting monitoring surveys of the Concho water snake, *N. h. paucimaculata* (Janecka *et al*. 2021a). At each site, the snake hosts were captured by hand or with the use of partially submerged minnow traps baited with Little Stinker catfish bait. Parasites were removed from the mouth, heat-killed under a cover slip and placed in 70% EtOH for use in the molecular analysis. Snakes were processed within 24 hours of capture, pit tagged and returned to their original location.

### 2.2 Microsatellite Development and Genotyping

Three individual parasites from three different hosts were submitted to the Sequencing and Genotyping Facility at the Cornell Life Sciences Core Laboratory Center (Ithaca, NY) to develop a microsatellite library following protocols described in Nali et al. (2014) (see also van Paridon et al. 2016; Detwiler and Criscione 2011). Potential microsatellite loci were screened in a subset of 6 individuals from 6 different hosts. DNA was extracted by placing the anterior portion of the worm in 200 uL of 5% chelex and 0.2 mg/ml of protinase K, incubated for 2 hours at 56°C and boiled at 100°C for 10 min. Only the anterior was used to avoid genotyping alleles of potential sperm donors that might be amplified from eggs or stored sperm in these hermaphroditic parasites. The M13 method was used for screening and subsequent genotyping. An 18-bp M13 tag (TGTAAAACGACGGCCAGT) was attached to the 5′ end of the forward primer (Schuelke 2000). A short tail sequence (GTTTCTT) was added to the 5′ end of the reverse primer to reduce polyadenylation (Brownstein et al. 1996). PCR amplification was performed in 10 µL reactions containing 3.1 µL ultrapure water, 5 µl 2X Qiagen Type-IT kit Master Mix, 0.16 µL fluorescent-labeled M13 primer (Applied Biosystems: FAM), 0.08 µL M13-labeled forward primer, 0.16 µL of 10 µM reverse primer and 1.5 µL of genomic DNA. The thermocycler profile was 94 °C for 5 minutes, 31 cycles of 94 °C for 30 seconds, 56 °C for 45 seconds, 65 °C for 45 seconds, followed by nine cycles of 94 °C for 30 seconds, 53 °C for 45 seconds 65 °C for 45 seconds and extension at 65 °C for 10 minutes. PCR-product was visualized on a 2% agarose gel run in 0.5X TBE buffer at 95 °C for 45 minutes. Reactions that yielded discrete bands in the expected product size range were sent to the DNA Analysis Facility on Science Hill at Yale University (New Haven, CT, USA) and visualized on a 3730xl 96-capillary Genetic Analyzer with 500-LIZ size standard. Samples were genotyped on GeneMarker 2.6.4. Loci were included in the final panel if there was allelic variation, if alleles could be scored without ambiguity, and alleles were reproducible in repeated samples. The final panel consisted of 11 novel loci (Table 1). Individual parasites were sampled across multiple hosts in a given location to provide an overall representative estimate of overall parasite population diversity (see Table S1 for genotyped sample sizes per host). There were 28 individuals for whom four or more loci failed to amplify even after re-amplification was attempted. These individuals were removed from the data set as they likely indicate poor extraction quality.

**TABLE 1.** Microsatellite diversity of *R. aniarum* populations at 13 loci from 13 collection sites on the Colorado and Concho Rivers and 2 tributary locations after clonemates were identified and reduced to a single representative individual.

| Population | $N^a$ | $NH^b$ | $A_R^c$ | $H_o^d$ | $H_s^e$ | $F_{IS}^f$ | $g^g$ | $P^h$ |
| --- | --- | --- | --- | --- | --- | --- | --- | --- |
| Upper Colorado River |  |  |  |  |  |  |  |  |
| Hwy 277 | 36 | 13 | 8.18 | 0.712 | 0.810 | **0.121** | 0.0092 | 0.166 |
| Elm Creek | 37 | 8 | 8.96 | 0.698 | 0.816 | **0.145** | 0.0277 | **0.004** |
| Hwy 83 | 32 | 11 | 8.87 | 0.733 | 0.819 | **0.106** | 0.0045 | 0.284 |
| Potter Falls | 41 | 9 | 8.99 | 0.687 | 0.818 | **0.106** | 0.0386 | **0.005** |
| 12 Mile Bridge | 26 | 6 | 9.05 | 0.664 | 0.817 | **0.188** | 0.0088 | 0.166 |
| Lower Colorado River |  |  |  |  |  |  |  |  |
| Freese Dam | 87 | 30 | 8.78 | 0.680 | 0.795 | **0.145** | 0.017 | **0.009** |
| Fishing Hole | 29 | 8 | 8.95 | 0.721 | 0.775 | 0.071 | -0.0048 | 0.76 |
| Hwy 283 | 38 | 19 | 9.22 | 0.785 | 0.789 | 0.006 | 0.0075 | 0.07 |
| River Bend Ranch | 42 | 12 | 9.80 | 0.773 | 0.813 | 0.05 | 0 | 0.51 |
| Concho River |  |  |  |  |  |  |  |  |
| FM 380 | 15 | 6 | 8.83 | 0.752 |  | 0.075 | 0.01 | 0.2 |
| Heatchen Ranch | 21 | 6 | 9.27 | 0.727 | 0.812 | 0.087 | 0.02 | 0.5 |
| Lipan Creek | 19 | 6 | 9.55 | 0.770 | 0.797 | 0.068 | 0.009 | 0.2 |
| Paint Rock | 68 | 11 | 9.40 | 0.747 | 0.826 | 0.059 | -0.006 | 0.9 |

<sup>a</sup> Number of individuals genotyped after clonemates were reduced to a single representative individual
<sup>b</sup> Number of hosts the parasites were taken from
<sup>c</sup> Allelic richness
<sup>d</sup> Observed heterozygosity
<sup>e</sup> Gene diversity
<sup>f</sup> Bolded $F_{IS}$ values indicate significant deviations from HWE ( $p < 0.05$ )
<sup>g</sup> $g^2$ is a measure of identity disequilibrium
<sup>h</sup> Significance values for $g^2$ . Significant $g^2$ values indicate inbreeding may be driving elevated $F_{IS}$ values.

### 2.3 Identification of Clonemates

Trematode parasites reproduce asexually in the intermediate host, but sexually in the definitive host. Hence, it is possible to find clonemates (genetically identical individuals due to the asexual reproduction) in the samples of adult trematodes. To determine if individuals with identical multilocus genotypes (MLGs) were the product of asexual reproduction, we used GENALEX 6.503 (Peakall & Smouse 2012) to calculate the *P_sex_* values for each MLG with a significance cut off of *P_sex_* <0.05. The assessment of clonemates was conducted by reach (UCR, LCR and CHR) because *P_sex_* could be driven down by substructure. No identical MLGs were found across reaches (only within). If the *P_sex_* value of two or more copies of a MLG at n = 2 is significant, then all copies of the MLG are considered to be the product of clonal reproduction. One representative of each clone was maintained in downstream all analyses to prevent inflation of population structure due to clonemates (Prugnolle *et al*. 2005; Criscione *et al*. 2011). These individuals were not included in any subsequent analysis.

### 2.4 Individual-based clustering analysis

Isolation by distance patterns can be strongly violated by including only a few diverged outlier populations, especially when the outlier populations fall near the center of the sampling range (Taylor *et al*. 2003). For this reason, we conducted an individual-based clustering analysis that did not depend on a priori delimitations of populations to determine if there were outlier populations within the data set. We used the model-based Bayesian clustering implemented in STRUCTURE 2.3.4 (Pritchard et al. 2000), which partitions individuals based on HWE and linkage equilibrium. The input parameters for STRUCTURE were uncorrelated allele frequencies and no admixture following the recommendations of Wang (2017) for uneven sample sizes. STRUCTURE was run with 1,000,000 iterations with a burn-in of 100,000 iterations for K (i.e., the number of possible clusters) values 1 to 13 with 10 replications of each possible K value.

### 2.5 Population-level analyses

#### Genetic diversity and disequilibrium tests

Tests for genetic diversity and disequilibrium were conducted for each collection site. Genotypic disequilibrium for pairs of microsatellite loci was tested using GENEPOP web 4.2 (Raymond and Rousset 1995). Statistical significance was determined using the log likelihood ratio statistic (G-test) with Markov chain parameters all set for 5000 dememorizations, batches and iterations per batch. Gene diversity (*H_S_*), the number of alleles per locus (An), and allelic richness (AR) (rarefied number to smallest sample size of N =16) were calculated in adegenet and ade4 (Dray & Dufour 2007; Jombart & Ahmed 2011). Estimates of *F_IS_* (which quantifies the proportional change in heterozygosity due to deviations in Hardy Weinberg equilibrium, HWE) were calculated in FSTAT 2.39 (Goudet 2001). Tests for Identity Disequilibrium (ID) were conducted in InbreedR with a bootstrap value of 1000 and p value from 10,000 permutations (Stoffel et al. 2016). Testing for the presence of identity disequilibrium tests for inbreeding independent of *F_IS_* and assists in determining if high *F_IS_* values are due to inbreeding (which could result from self-mating) or are artificially inflated due the presence of technical scoring errors (e.g., null alleles).

#### Straight-line distance, river distance and genetic diversity

The stream-drift hypothesis predicts a negative correlation between genetic diversity measures and distance relative to the most downstream collection site (i.e., genetic diversity increases going downstream). We tested for a correlation between river distance (relative to the most downstream network position) and allelic richness (*AR*), or gene diversity (*H_S_*). Gene diversity measures did not meet assumptions of normality and were arcsine square root transformed prior to the correlation analysis (Whelan *et al*. 2019). Linear regressions were done in R. Statistical tests for differences in average genetic diversity indices between locations of the river were examined using the non-parametric Kruskal-Wallis test and implemented in R using the dplyr and coin libraries. Definitive host species *(Nerodia* spp.) are potentially capable of both terrestrial and aquatic dispersal and so have the potential to transmit *R. aniarum* infections through out-of-network host movement (i.e., terrestrial travel). To determine which of these potential dispersal paths was most associated with the distribution of *R. aniarum* genetic diversity, we conducted linear regressions of straight-line distance (Euclidean distances) and river distance (distance between sites following network pathways) for each of the response variables (allelic richness and gene diversity). River distances between collection sites were determined in ARCGIS 10.5.1 (ESRI Inc.). Distances were measured from the most downstream location of one site to the most upstream point of the next site (Paz-Vinas *et al*. 2015). Network position was measured as distance from the most downstream site on the Colorado River (River Bend Ranch). Straight-line distance between the centers of each collection site were calculated in adegenet 2.1.1 (Jombart & Ahmed 2011).

#### Population structure and isolation by distance along network pathways

Pairwise tests of genetic differentiation and estimation of *F_ST_* among sites were calculated in adegenet (Jombart & Ahmed 2011). We also conducted a principal coordinate analysis (PCoA) as implemented in GENALEX 6.502 (Peakall and Smouse 2012).

Mantel tests for correlation analyses between river distance or straight-line distance and pairwise linearized *F_ST_* (*F_ST_* /(1-*F_ST_*)values were calculated in adegenet 2.1.1 ^(^Jombart & Ahmed 2011^)^. Euclidean straight-line distance for Mantel tests were log transformed following the recommendations for two dimensional habitats (Rousset 1997). The STRUCTURE and PCoA analyses indicated that strong genetic structure associated with the Freese Dam location, which is centrally located within the river bifurcation. For this reason, the Freese Dam location was excluded from tests for IBD. Additionally, because we interested in linear networks along the mainstem of the river, the two tributary locations were also excluded from IBD analyses. Mantel tests were thus performed for a data set which included 10 populations on the mainstem of the three reaches of the river. Lastly, three subset Mantel tests (excluding the outlier at Freese Dam and the two tributary populations) were conducted for three river distance paths: 1) a ‘linear’ pathway using populations from UCR to LCR, 2) a ‘linear’ pathway using populations from CHR to LCR, and 3) a bifurcated pathway populations from UCR to CHR (Figure 1 but also see insets in Figure 4). A Bonferroni correction for multiple tests was applied to the results.

## 3. Results

### 3.1 Parasite Collections and Clonemate Identification

A total of 267 water snakes were captured and 1445 *R. aniarum* individuals were collected during the 2013-2015 sampling seasons from the 13 collection sites. Eleven microsatellite markers were amplified for 583 *R. aniarum* individuals collected at 13 sites along the Colorado and Concho Rivers and the two tributary locations. Individuals with missing data indicative of low-quality DNA extractions were removed, leaving 555 individuals with complete multilocus genotypes. There were no matching MLGs between any of the 13 sampled locations. Identical MLGs were only found within locations (Table S2). Clonemates (i.e., *P_sex_* < 0.05) were identified primarily within the same host individual but there were some clonemates taken from two hosts captured within the same location (Table S2, see Elm Creek and Potter Falls for example). Clonemates from the 555 individuals were reduced to a single representative of each clone for subsequent analyses. The remaining data set included in the analysis included 491 *R. aniarum* individuals from the 13 collection locations (Table 1).

### 3.2 Individual-based clustering analysis

The Bayesian clustering analysis revealed population subdivision associated with the population located immediately below Freese Dam (Figure 3a). There is also moderate but visible differentiation between all the populations on the UCR relative to the CHR and LCRs. There is no visible distinction in genetic clusters between the CHR and LCR. The mean of the estimated Ln probability of the data suggests that 6 is the most likely number of genetic clusters (Figure S1). These clusters are not associated with geographic locations or with host species (Figure 3). No additional population substructure is visible among the three reaches of the river.

**FIGURE 3.**
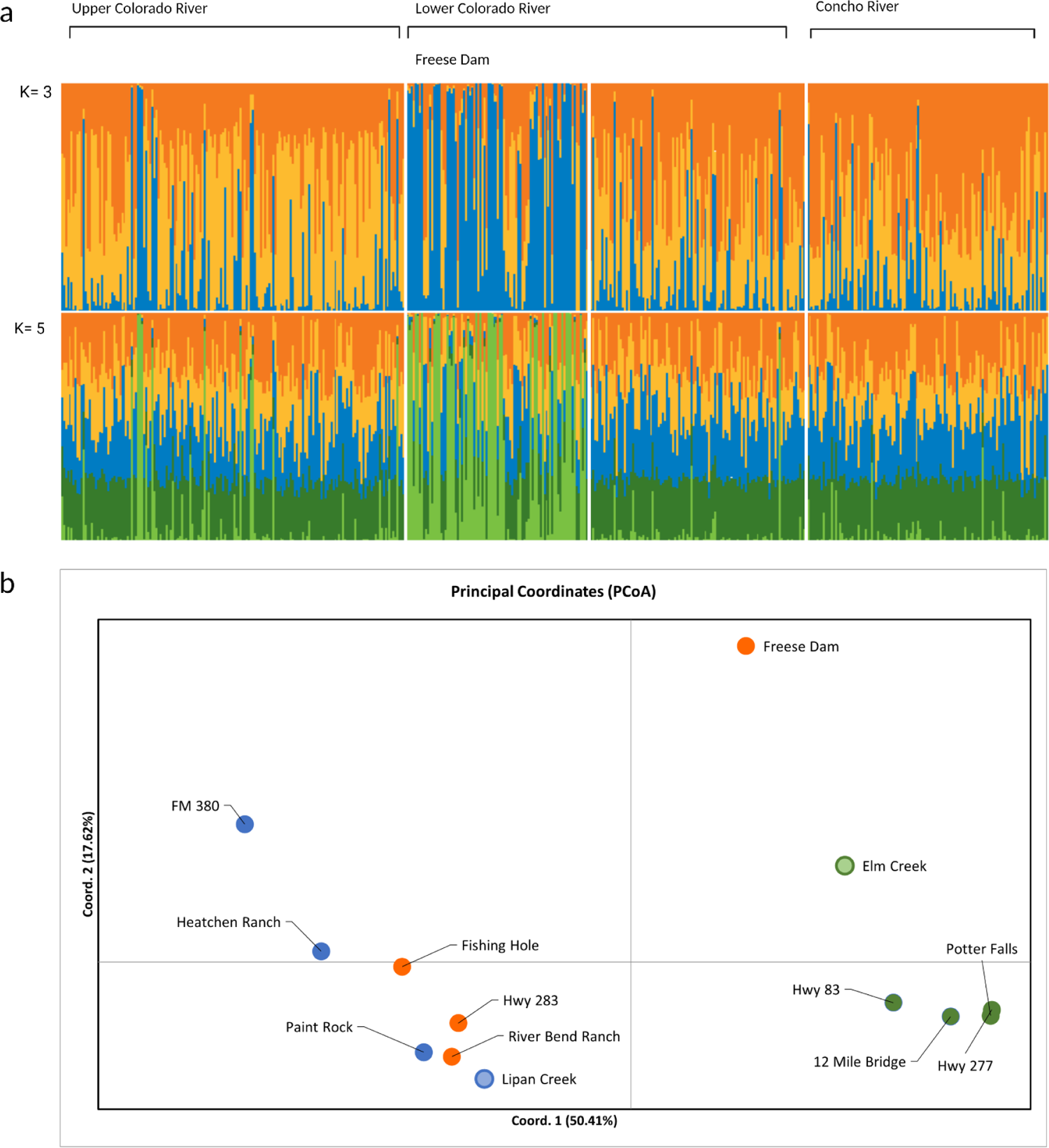
Results from the Baysian clutering analysis of *R. aniarum* with no population as prior assumed. Individuals are arranged from most upstream to most downstream location on the Colorado River and then most upstream to most downstream on the Concho River. Each individual is represented by a vertical bar divided colors which represent the diffeent genetic clusters. The length of the color is proportional to the to the probability of assignment to each cluster. The Freese Dam population is located immediately below O.H. Ivie reserviour but is visibly distinct,despite its cetral geographic location

### 3.3 Population-level analyses

#### Genetic diversity and disequilibrium tests

The number of loci in each population that exhibited significant deviations from HWE ranged from 0 to 6 (6 from Potter Falls). The loci that exhibited significant deviations varied between collection sites and no markers exhibited significant deviations from HWE at all locations. Multi-locus *F_IS_* values ranged from 0.006 (Hwy 283) to 0.188 (12 Mile Bridge) (Table 1). Sites located on the UCR and directly downstream of Freese Dam all had *F_IS_* values significantly greater than 0. However, *F_IS_* values were not significantly greater than 0 for the remainder of sites on the LCR and for no sites on the CHR.

We found significant levels of identity disequilibrium in three populations, Elm Creek, 12 Mile Bridge, and Freese Dam (Table 1), suggesting that the presence of null alleles may be inflating *F_IS_* values in Hwy 277, Hwy 83, and Potter Falls. No other sites along the CHR and the remainder of LCR had significant levels of identity disequilibrium.

Linkage disequilibrium tests were performed at the level of collection sites. For 11 microsatellite markers there are 55 pairwise comparisons between loci. At an alpha of 0.05, one would expect 2.75 loci per population to be in LD by chance alone. Among the 13 populations, 0 to 3 pairs of loci tested significant for LD (only 12 Mile Bridge had 3 significant pairs). No pairs of loci exhibited LD consistently among locations. Hence, we found no strong evidence for LD.

### Straight-line distance, river distance and genetic diversity

Linear regressions between both gene diversity or allelic richness, and river distance or straight-line distance were not significantly different among sites (Table 2). Levels of gene diversity were high for all populations (Table 1), and average across loci *Hs* was not significantly different among the 13 locations based on the Kruskal Wallis test (χ ^2^=2.183, p=0.9991, df=12). Allelic richness was similarly high and not significantly different among groups (χ ^2^=1.683, p=0.9998, df=12).

**TABLE 2.** Results of the linear regressions to examine the relationship between genetic distance and distance upstream (river distance, km) and straight-line distance on average genetic diversity metrics per site for *R. aniarum*. Gene diversity (*H*_S_) is arcsine square root transformed prior to use in the correlation analysis.

|  | River Distance |  | Straight-line Distance |  |
| --- | --- | --- | --- | --- |
| | $R^2$ | $p$ | $R^2$ | $p$ |
| Gene Diversity ($H_S$) | -0.003 | 0.4924 | -0.0032 | 0.465 |
| Allelic Richness | -0.0049 | 0.583 | -0.0049 | 0.588 |

### Population structure and isolation by distance along network pathways

Pairwise comparisons of *F_ST_* values between sites were overall relatively low (Table 3). Wier and Cockerham (1984) theta estimates however indicated significant population subdivision with increasing geographic distance among sites except for sites on the LCR and CHR, which did not indicate significant population subdivision despite being located on separate reaches of the river and with geographic river distances of 50 to 150 km among sites (see the comparison between Hwy 283 and Heatchen Ranch as an example). Freese Dam, the population located immediately downstream from O.H. Ivie Reservoir, which is at the confluence of the Colorado and Concho Rivers, was also significantly differentiated from all other sampling locations, including Fishing Hole which is only 10 river km downstream from Freese Dam.

**TABLE 3.** Pairwise measure of *F_ST_* below the diagonal. Significance values are given above the diagonal based on Weir and Cockerham (1984) theta estimate based on 999 permutations.

| Population | Hwy<br>277 | Elm<br>Creek | Hwy<br>83 | Potter<br>Falls | 12<br>Mile<br>Bridge | Freese<br>Dam | Fishing<br>Hole | Hwy<br>283 | River<br>Bend<br>Ranch | FM<br>380 | Heatchen<br>Ranch | Lipan<br>Creek | Paint<br>Rock |
| --- | --- | --- | --- | --- | --- | --- | --- | --- | --- | --- | --- | --- | --- |
| Hwy 277 | - | 0.014 | **0.009** | **0.001** | **0.013** | **0.001** | **0.001** | **0.001** | **0.001** | **0.001** | **0.001** | **0.001** | **0.001** |
| Elm Creek | 0.006 | - | 0.183 | **0.006** | 0.124 | **0.001** | **0.001** | **0.001** | **0.001** | **0.001** | **0.001** | **0.001** | **0.001** |
| Hwy 83 | 0.008 | 0.003 | - | 0.441 | 0.462 | **0.001** | **0.001** | **0.001** | **0.002** | **0.001** | **0.002** | **0.014** | **0.001** |
| Potter Falls | 0.011 | 0.008 | 0.000 | - | 0.159 | **0.001** | **0.001** | **0.001** | **0.001** | **0.001** | **0.001** | **0.002** | **0.001** |
| 12 Mile<br>Bridge | 0.009 | 0.004 | 0.000 | 0.003 | - | **0.001** | **0.001** | **0.002** | **0.001** | **0.001** | **0.001** | **0.002** | **0.001** |
| Freese Dam | 0.024 | 0.012 | 0.017 | 0.022 | 0.019 | - | **0.001** | **0.001** | **0.001** | **0.002** | **0.001** | **0.001** | **0.001** |
| Fishing Hole | 0.030 | 0.020 | 0.017 | 0.026 | 0.024 | 0.025 | - | **0.014** | **0.002** | **0.007** | 0.068 | 0.021 | **0.004** |
| Hwy 283 | 0.022 | 0.016 | 0.014 | 0.019 | 0.016 | 0.019 | 0.008 | - | 0.423 | **0.012** | 0.104 | 0.055 | 0.255 |
| River Bend<br>Ranch | 0.021 | 0.014 | 0.012 | 0.019 | 0.015 | 0.023 | 0.013 | 0.000 | - | 0.025 | 0.063 | 0.406 | 0.283 |
| FM 380 | 0.040 | 0.019 | 0.027 | 0.034 | 0.034 | 0.026 | 0.017 | 0.015 | 0.010 | - | 0.230 | 0.011 | **0.009** |
| Heatchen<br>Ranch | 0.028 | 0.015 | 0.017 | 0.028 | 0.020 | 0.022 | 0.006 | 0.005 | 0.005 | 0.004 | - | 0.199 | 0.254 |
| Lipan Creek | 0.022 | 0.015 | 0.011 | 0.017 | 0.016 | 0.025 | 0.011 | 0.007 | 0.000 | 0.017 | 0.004 | - | **0.020** |
| Paint Rock | 0.022 | 0.017 | 0.015 | 0.022 | 0.016 | 0.025 | 0.010 | 0.001 | 0.001 | 0.012 | 0.002 | 0.008 | - |

The PCoA analysis separated the UCR from the LCR and CHR along the x-axis (50.41%) (Figure 3b). Sites from the CHR and LCR formed a cluster primarily in the lower left quadrant. The exception was Freese Dam, which fell away from all other sites. These results of the PCoA were largely concordant with the pairwise *F_ST_* analysis and the STRUCTURE analysis, which suggest low population subdivision between the LCR and CHR reaches.

Mantel tests for IBD for the 10 mainstem populations indicate significant isolation by river distance but not by straight line-distance after the Bonferoni correction (Figure 4, Figure S2 and Table 4). It should be noted that river distance and straight-line distances along network pathways between the UCR and LCR were very similar. For the linear pathway along the UCR and CHR, there was no significant IBD for river distance after the Bonferroni correction, however there was IBD by straight-line distance and a higher R^2^ value. There was no significant IBD for either straight-line or river distance along the Linear pathway between the LCR and CHR rivers. The most significant test for IBD and highest R^2^ value was for the bifurcated network river pathway between the UCR and CHR, indicating that there was a highly significant correlation between the pairwise linearized *F_ST_* matrix and river distance matrix. In contrast there was no IDB detected for the out of network straight-line distances between the two forked reaches of the river (Figure 4, Figure S2 and Table 4).

**FIGURE 4.**
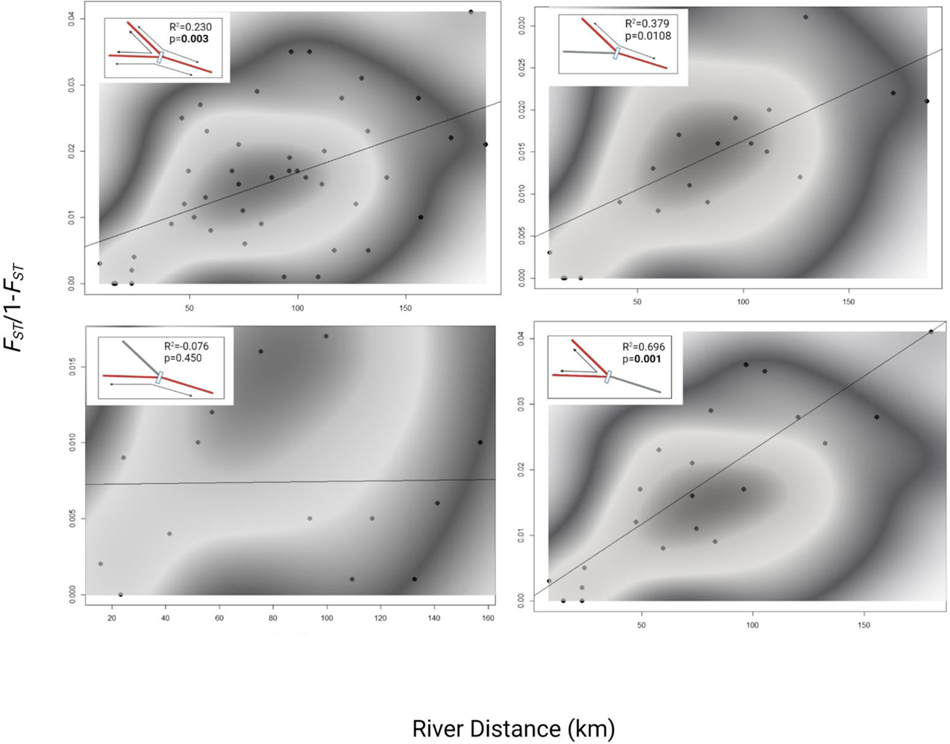
Correlation between matrices of geographic distance and population linearized pairwise distances among sites for sites along linear pathways within the Colorado and Concho River (Mantel test values for significance given in figure, r values in Table 4). Heat map is overlaid to show local density data. Analysis excludes outlier (Freese Dam) and tributary locations.

**TABLE 4.** Results for the Mantel Tests for Linearized *F_ST_* (*F_ST_* /1-*F_ST_*) and straight-line Euclidean distance (log transformed). After Bonferroni correction for multiple tests, significance is assigned at 0.006

| Network Pathway | R<sup>2</sup> | River Distance<br>(km) P | R<sup>2</sup> | Straight Line Distance<br>(log km) P |
| --- | --- | --- | --- | --- |
| Mainstem<br>populations (10) | 0.230 | **0.003** | 0.102 | 0.018 |
| Upper and Lower<br>Colorado River | 0.379 | 0.0108 | 0.484 | **0.005** |
| Concho River and<br>Lower Colorado<br>River | -0.076 | 0.450 | 0.097 | 0.037 |
| Upper Colorado<br>River and Concho<br>River | 0.690 | **0.001** | 0.129 | 0.04 |

## 4. Discussion

### Is stream drift shaping R. aniarum genetic diversity?

In trematode parasites with complex life cycles, parasite dispersal and genetic structure is frequently determined by the host with the greatest dispersal (Jarne & Théron 2001; Prugnolle *et al*. 2005; Louhi *et al*. 2010). In this study, we examined the relative influence of unidirectional stream drift, host dispersal and dendritic branching in shaping the genetic diversity and population structure of a parasite infecting semi-aquatic snakes. Unidirectional river currents would drive down-stream parasite dispersal, whereas upstream gene flow must be facilitated by upstream host dispersal. *Renifer aniarum* exhibited high levels of genetic diversity that were not significantly different among sites or among the three reaches of the river. Hence, genetic diversity did not decrease with distance upstream as would be predicted if unidirectional stream drift was a significant process shaping the distribution of parasite genetic diversity. Thus, neither position within the river network nor reach influenced the distribution of genetic diversity. The distribution of *R. aniarum* genetic diversity is high and similar among locations regardless of relative position along this river system, suggesting similar effective population sizes of the parasite among the reaches of this river system.

Balsco Costa et al. (2012) posited that parasites of hosts with dispersal abilities that can counter unidirectional stream drift will not exhibit a decrease in genetic diversity with distance upstream. Most members of the genus *Nerodia* maintain relatively large home range sizes (2-8 ha), although there is considerable variation in home range size between sexes, species and locations (Pattishall & Cundall 2008; Camper 2009; Camper & Chick 2010). Area use of the snake hosts scales up with host size, with larger snakes moving long distances during the active months in pursuit of food and mating opportunities (Todd & Nowakowski 2021). Consistent with these behaviors, previous work with nuclear markers found no evidence of population structure for *N. erythrogaster* on the Colorado and Concho Rivers (Rodriguez *et al*. 2012). Thus, these host patterns along with our results of the parasite’s genetic diversity are consistent with the hypothesis proposed by Balsco Costa et al. (2012).

### Does the DEN shape R. aniarum population structure?

We predicted that if the DEN was a major determinate of *R. aniarum* population structure, we would find population subdivision associated with the three reaches of the river and patterns of isolation by distance along bifurcating network pathways. We found no strong patterns of IBD along linear network pathways, either along the UCR-LCR or LCR and CHR after applying a Bonferroni correction for multiple tests. In contrast, we found strong patterns of isolation by distance associated with the network bifurcations, with the highest R^2^ values associated only driven by pathways between the UCR and CHR. We also found high *F_IS_* values in the UCR. Heterozygote deficiencies in natural populations can be generated by null alleles, the Wahlund effect, self-fertilization, and positive associative mating (Waples 2015).

Microsatellite null alleles are loci that fail to amplify sufficiently for detection during genotyping. Estimates of identity disequilibrium suggest that the presence of null alleles is the significant contributing factor for three out of these six populations, while inbreeding and selfing are the dominate mechanism in the remaining three. Populations from the LCR (except Freese Dam) and the CHR river has both *F_IS_* values approaching zero and no evidence of identity disequilibrium. The geographic restriction of inbreeding and null alleles to the UCR and Freese Dam suggests null alleles may be circulating in the UCR but absent from the LCR and CHR, making the pattern biologically relevant.

We also predicted that the DEN would generate patterns of population subdivision associated with the three reaches of the river. However, our results revealed more subtle patterns of population subdivision, consistent with IBD. Pairwise *F_ST_* values were relatively low but increased and became significant with increasing distance among sites. Subpopulations located at the most distal, upstream locations of both the UCR, CHR and below Freese Dam were significantly differentiated from all other sites. The results of the PCoA and the STRUCTURE analysis emphasize some geographic differentiation for sites on the UCR relative to the LCR and CHR. The results of the STRUCTURE analysis also identified strong population associated with Freese Dam. Isolation by distance patterns have been previously determined to reduce the efficacy of the program STRUCTURE and lead to false identification of genetic clusters (Perez *et al*. 2018). It is likely that the IBD present within the data set influenced the programs identification of a best K values of 5, as suggested by the mean Ln probability of the data. *Do hosts circumvent the barriers to aquatic systems?*

We predicted that if host-driven parasite dispersal circumvented the barriers to aquatic systems, then we would find no population structure and no patterns of isolation by distance due to transmission of parasites by the primary definitive host in the system, *N. erythorgaster*. We found no correlation between Euclidean distance and pairwise genetic distance, but a highly significant correlation for river distance within the bifurcating network. Thus, the comparison of terrestrial versus within network dispersal pathways indicates that while water snakes may hypothetically be capable of terrestrially facilitating parasite gene flow between network branches, actual parasite dispersal is dominated primarily by dispersal within the river network. Our results indicate that the structural feature of river bifurcation and inherent variation in connectivity between river branches shape *R. aniarum* population structure and isolation by distance even when host dispersal is high. However, our results to do not rule out the possibility of short distance out of network dispersal by water snake hosts facilitating *R. aniarum* transmission. *Nerodia erytrhogaster* is the most terrestrial of all watersnake species (Gibbons and Dorcus 2004). The correlation between straight line distance and genetic distance along the UCR and LCR linear pathway was more significant and had a higher *R^2^* value than river distance along this same gradient. While both distances are similar, straight line distance movement between these rivers is a shorter pathlength. These results suggest the possibility that watersnake terrestrial dispersal along linear pathways may also be important in *R. aniarum* transmission within the river. Short distance terrestrial dispersal along linear networks but not between branches has been documented in other freshwater semi-aquatic vertebrates, including Eurasian otters (Carranza et al. 2012), turtles (Ciofi et al. 2016; McGaugh 2012), and salamanders (Mullen et al. 2010; Olson and Kluber 2014). Our research is the first to examine how this variation in terrestrial dispersal (short-distance linear but not between branches) events may influence parasite dispersal. Further work is required to determine how additional landscape features such as variation in habitat permeability may play a role in facilitating semi-aquatic host parasite transmission in riverine ecosystems.

### Reservoir Effects

The Freese Dam location was an outlier within this river system. We did not make any a priori hypotheses regarding the effects of O.H. Ivie reservoir on *R. aniarum* population structure. *Renifer aniarum* individuals collected from the Freese Dam site were consistently diverged from all other sampling locations independent of geographic distance. It is of note that this same pattern of population subdivision associated with the reservoir was also found for the recently delisted host, *N. harteri paucimaculata*, whose post-delisting monitoring was conducted concurrently with this study (Janecka et al. 2021). We do not have a current biological explanation for the concordance of the results for both this host and the parasite, however two organisms whose data independently identifies the same pattern warrants further consideration.

The Freese Dam collection site is located immediately downstream from O.H. Ivie reservoir, near the confluence of the Colorado and Concho Rivers. Construction was completed on O.H. Ivie reservoir in 1989, approximately 30 years prior to when these samples were collected. The construction of impoundments in rivers are one of the most widespread alterations to river ecosystems (Theime et al. 2020; Thieme et al. 2021;). Reservoirs have been shown to alter the metapopulation structure and dynamics of small fish species (Falke & Gido 2006), generated patterns of decreased genetic diversity upstream (Dehais *et al*. 2010), and generate population subdivision in rivers fragmented by reservoirs (Sotola *et al*. 2017). Atkinson and Bartholomew (2010) studied the population structure of the myxozoan parasite *Ceratomyxa shast*a in several species of salmonid fish in the Kalmath River 88 using the mitochondrial ITS-1 marker. Dams have separated the upper, middle and lower basins of the river for approximately 80 years. Atkinson and Bartholomew (2010) found the population structure of *Ceratomyxa shasta* (Myxozoa) was highly structured spatially between reaches of the river separated by dams and between host species. Distinct *C. shasta* genotypes were restricted to river basins isolated between dams, indicating that the dam was likely responsible for the isolation of the parasite between reaches. Pettersen et al. (2015) also state that the presence of reservoirs act as a barrier to host-parasite dispersal with their river network but did not address the length of time these barriers have been in place. Our results did not find evidence of long-term disruptions in population connectivity between other *R. aniarum* populations, possibly driven by large metapopulation sizes and insufficient time since construction which may mitigate the effects of the reservoir on connectivity along the Concho-Lower Colorado reach of the river.

It is currently unclear what may be driving the population subdivision immediately downstream of Freese Dam. It is possible that smaller effective population sizes and lack of migrants from upstream may have increased the influence of drift at this site relative to the effects of migration. Alternatively, one or more genetic bottlenecks immediately below the reservoir followed by subsequent recolonization by colonists dominated by a particular genotype could generate local population subdivision. It is of note that Zhang et al. (2010) found a similar population subdivision in the largemouth bronze gudgeon (*Coreius guichenoti*) immediately below the Three Gorges Dam on the Yangtze River while other sites maintained high connectivity. However, it is possible the presence of O.H Ivie reservoir may have impacted *R. aniarum* transmission in other ways. The presence of dams has been documented to significantly alter the transmission of human blood flukes, *Schistosoma mansoni*, by reducing salinity and thus enabling colonization by the freshwater snail host, *Biomphalaria pfeifferi* (Campbell *et al*. 2010). Diakite et al. (2017) reported increases in snail abundance and changes in community composition in response to the construction of new dams due to changes in habitat stability from flood control. Increased habitat predictability can also alter snail life history characteristics such as fecundity and survivorship, characteristics that can also influence trematode transmission (Diamond 1982; Charbonnel *et al*. 2002). Characteristics of the river immediately below O.H. Ivie reservoir are distinct from other locations along the Colorado and Concho Rivers, with dense aquatic vegetation and wide marshy areas adjacent to the riverbed and significant accumulation of fine-particle silt, suggesting greater habitat stability at this collection site (Janecka, personal observation). It is possible that the habitat alteration may have in turn altered transmission between the first and or second intermediate host, changes in intermediate host density or community composition. However, while the significant population subdivision below Freese Dam is interesting and warrants further investigation. Replication of large-scale studies of the effects of reservoirs on parasite population subdivision are required to further understand how impoundments may alter parasite dispersal in riverine systems.

## Conclusions and Conservation Implications

Our understanding of the effects of the unique characteristics of riverine habitats in shaping the ecology and evolution of their inhabitants is a rapidly expanding field that is still based more in theoretical and simulation data sets rather than empirical work for parasites that infect riverine hosts. We found the strongest population structure immediately below the reservoir. Patterns of isolation by distance were not uniform along network pathways, with significant IBD and genetic differentiation associated with the UCR, despite similar path network path lengths along the CHR-LCR pathway. The comparison of terrestrial versus within network dispersal pathways indicates that while water snakes may be capable of terrestrially facilitating parasite gene flow between network branches, actual parasite dispersal is dominated primarily by dispersal within bifurcations and network short terrestrial dispersal may be important along linear networks.

Our results also have direct conservation implications within the Colorado and Concho Rivers. Snake fungal disease (*Ophidiomyces ophiodiicola*) has recently been reported infecting the watersnake community in the Brazos River, including the endemic Brazos watersnake (*Nerodia harteri harteri*), a close relative of the recently de-listed endemic Concho watersnake (*N. h. paucimaculata*) included in this study (Harding et al. 2022). Both *N. harteri* subspecies are highly aquatic and rarely found more than a meter from water and exhibit short distance dispersal (Gibbons and Dorcus 2004; Janecka et al. 2021). Snake fungal disease causes morbidity in the form of pustules, lesions and crusty scales and causes morbidity and mortality in some species. We did not detect visible infection of SFD during the course of this study (Janecka personal obsv). While infection by snake fungal disease also circulates in terrestrial habitats, our results suggest the possibility that watersnake-to-watersnake transmission may be highest along linear pathways between the UCR and CHR, should this pathogen be introduced to the system.

By explicitly addressing the effects of dendritic branching on parasite dispersal within river networks, our results highlight that the complexity natural host-parasite systems necessitate future empirical data from natural systems. This foundational knowledge is required to capture the complexity of parasite transmission in river ecosystems within predictive models of both disease spread and parasite adaptive potential within rivers (Thrush et al. 2011; Chiu et al. 2020; Ma et al. 2020).

