## Supplemental materials for "Parasite gene flow in riverine habitats: ascertaining the roles of stream drift, river bifurcations and host dispersal"

**Table S1.** The 11 microsatellite loci developed for *R. aniarum* in this study.

| Primer Name | Primer Sequence (5' to 3') | Repeat Motif | $T_a$(ºC) | Size Range (bp) |
|---|---|---|---|---|
| **ReAn 627** | F: TGTAAAACGACGGCCAGT | AGC | 56 | 347-426 |
| | R: GAGACTTCAAGCCTCGAACTG | | | |
| **ReAn 1082** | F: GGCTAACCCAGTTTCGAAGC | ACT | 56 | 244-378 |
| | R: TGAGTCGGCATTAGTTGATTGG | | | |
| **ReAn 5773** | F: ACTGCCAACACTCGATTGATC | ACAG | 56 | 170-304 |
| | R: ACGTACAATAAATGCAGCGC | | | |
| **ReAn 8483** | F: TTCAACCACACTTCGTTCGC | AC | 56 | 155-193 |
| | R: AGTTCAACTGATCAACGACGAG | | | |
| **ReAn 3858** | F: GGAGCTTGCCACAACCAC | AAC | 56 | 155-260 |
| | R: GACTCGGCGTTCTAAAGTGC | | | |
| **ReAn 2341** | F: ACCCATGCAACCAAGATGAG | AG | 56 | 380-406 |
| | R: GGTACTCTCCTGGGATCGTG | | | |
| **ReAn 5592** | F: TGACCGCAGATTTGACCAATG | ATC | 56 | 298-366 |

| | | | | |
|---|---|---|---|---|
| | R: GGTCTACGTGCTTGTGTTCG | | | |
| **ReAn 990** | F: ACCCGCTTGTCATATTCAGTG | AAC | 56 | 169-202 |
| | R: TGAACCCACAATTTCGCTGG | | | |
| **ReAn 1248** | F: TCGGAGAACAAACCACCAC | AAG | 56 | 395-464 |
| | R: AACTTGAAGCAAAGGTGGCC | | | |
| **ReAn 2450** | F: CTAAAGCGTCTCGTTCCTTGATTC | ACAT | 56 | 135-173 |
| | R: GTCACAATGGAGTTTCAAATGTCG | | | |
| **ReAn 1065** | F: AGTGCCTGTTCGCTGATTTC | AAG | 56 | 161-215 |
| | R: CTGTGGAAAGTCGTTGGTATTG | | | |

**Table S2.** The distribtion of identical mulilocus genotypes among hosts whose $P_{sex}$ values were significant ($P_{sex}$ <0.05). Host IDs are used for categorization within the table but are not the same individual snakes across sites. Groups of clone mates with the same MLG ID were reduced to a single representative individual for all other analyses.

| Site | Hwy 277 | | Elm Creek | | Hwy 83 | | Potter Falls | | 12 Mile Bridge | | Freese Dam | | Fishing Hole | | Hwy 283 | | River Bend Ranch | | FM 380 | | Heatchen Ranch | | Lipan Creek | | Paint Rock | |
|---|---|---|---|---|---|---|---|---|---|---|---|---|---|---|---|---|---|---|---|---|---|---|---|---|---|---|
| Host ID[a] | N[b] | MLG ID[c] | N | MLG ID | N | MLG ID | N | MLG ID | N | MLG ID | N | MLG ID | N | MLG ID | N | MLG ID | | MLG ID | N | MLG ID | N | MLG ID | N | MLG ID | N | MLG ID |
| 1 | 1 | | 7 | | 2 | | 2 | | 7 | | 2 | | 2 | | 1 | | 2 | | 3 | S(2) | 3 | | 4 | W(2) | 3 | |
| 2 | 3 | | 7 | | 2 | | 8 | E(1) F(2) | 2 | | 1 | | 4 | M (2) | 4 | O(2) | 2 | | 2 | T(2) | 4 | | 6 | | 3 | |
| 3 | 1 | | 3 | B(2) | 3 | | 8 | | 6 | | 1 | | 6 | | 1 | | 4 | | 3 | U(2) | 2 | | 2 | | 5 | |
| 4 | 5 | | 6 | C(2) | 3 | | 11 | | 5 | | 3 | | 4 | | 1 | | 4 | | 4 | | 5 | | 2 | | 2 | |
| 5 | 5 | | 1 | | 3 | | 2 | | 5 | | 1 | | 4 | N (3) | 2 | | 3 | | 4 | V(2) | 3 | | 1 | | 3 | |
| 6 | 3 | | 7 | | 3 | | 9 | E(2) F(1) G(2) | 1 | | 3 | I(1) | 4 | | 4 | | 4 | Q (1) | 4 | | 4 | | 5 | | 3 | |
| 7 | 2 | | 6 | B(2) | 5 | | 1 | | | | 1 | | 4 | | 2 | | 4 | | | | | | | | 3 | |
| 8 | 5 | | 4 | | 3 | | 4 | H(2) | | | 1 | | 4 | | 1 | | 1 | | | | | | | | 26 | X(3) Y(3) Z(3) AA(3) BB(2) |
| 9 | 2 | | | | 1 | | 2 | | | | 1 | | | | 1 | | 4 | | | | | | | | 4 | CC(2) |
| 10 | 2 | A (2) | | | 4 | D(2) | | | | | 2 | | | | 3 | | 5 | Q(1) | | | | | | | 14 | DD(2) EE(3) |
| 11 | 1 | | | | 4 | | | | | | 1 | | | | 2 | | 9 | Q(1) R(2) | | | | | | | 21 | FF(2) GG(2) HH(2) FF(2) II(2) |
| 12 | 5 | | | | | | | | | | 1 | | | | 3 | | 3 | | | | | | | | | |
| 13 | 2 | | | | | | | | | | 3 | | | | 2 | | | | | | | | | | | |
| 14 | | | | | | | | | | | 5 | | | | 2 | P(2) | | | | | | | | | | |
| 15 | | | | | | | | | | | 1 | | | | 4 | | | | | | | | | | | |

| | | | | | | | | | | | | | | |
|---|---|---|---|---|---|---|---|---|---|---|---|---|---|---|
| **16** | | | | | | | | | | 3 | | | | 2 |
| **17** | | | | | | | | | | 12 | | | | 3 |
| **18** | | | | | | | | | | 5 | J(2) | | | 1 |
| **19** | | | | | | | | | | 1 | | | | 1 |
| **20** | | | | | | | | | | 1 | | | | |
| **21** | | | | | | | | | | 3 | | | | |
| **22** | | | | | | | | | | 16 | K(2) | | | |
| **23** | | | | | | | | | | 7 | L(2)<br>I(1) | | | |
| **24** | | | | | | | | | | 1 | | | | |
| **25** | | | | | | | | | | 1 | | | | |
| **26** | | | | | | | | | | 1 | | | | |
| **27** | | | | | | | | | | 13 | | | | |
| **28** | | | | | | | | | | 2 | | | | |

[a] The host from which the parasites were collected. Host ID's are not the same individuals across sites.

[b] The number of *R. aniarum* individuals genotyped from that host after individuals with missing data were removed

[c]MLG ID Indicates individuals with identical MLGs.  These individuals were reduced to a single representative within the data set.

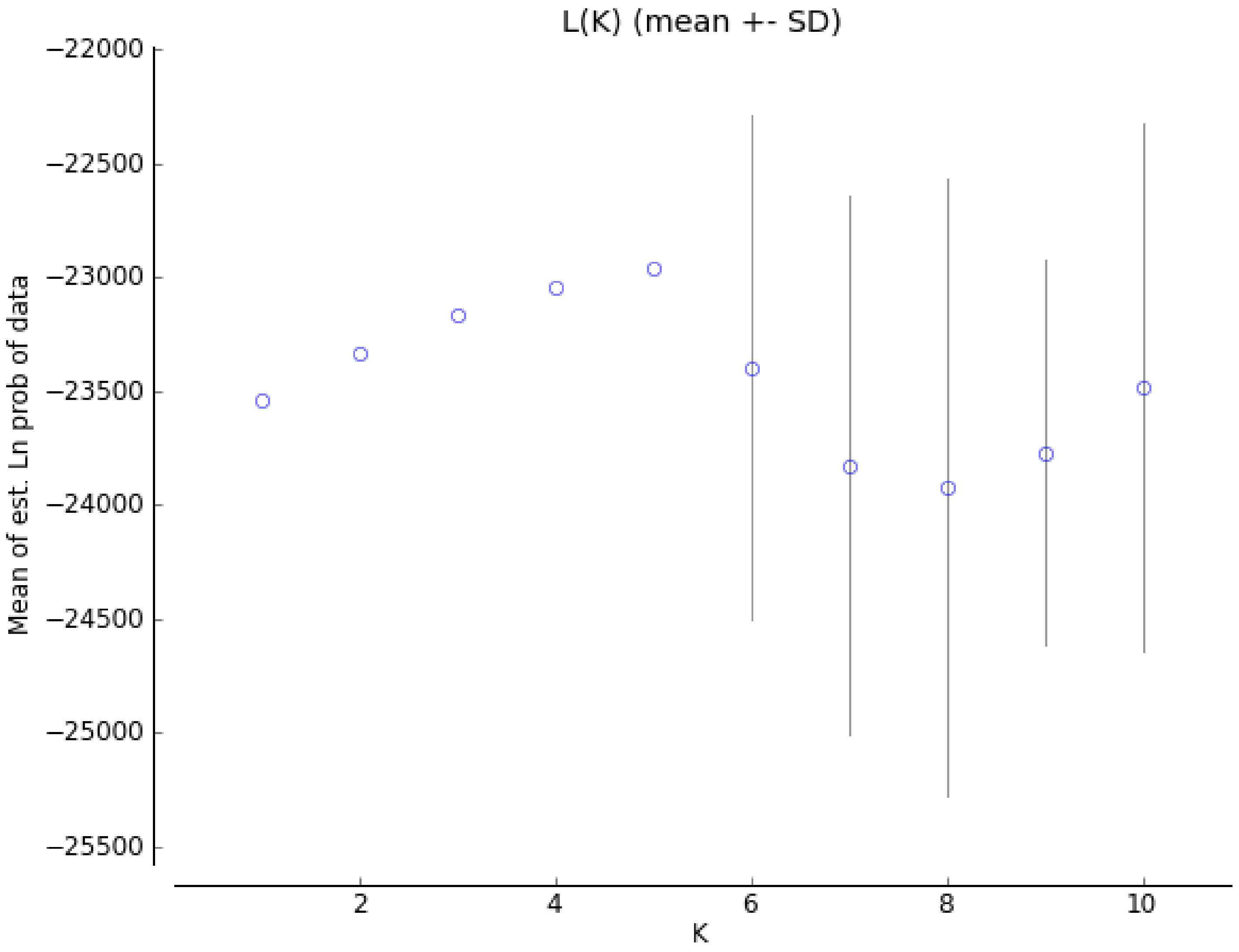

**Figure S1.** Mean of the estimated Ln probability of the data for each K and standard deviations from the STRUCTURE analysis.

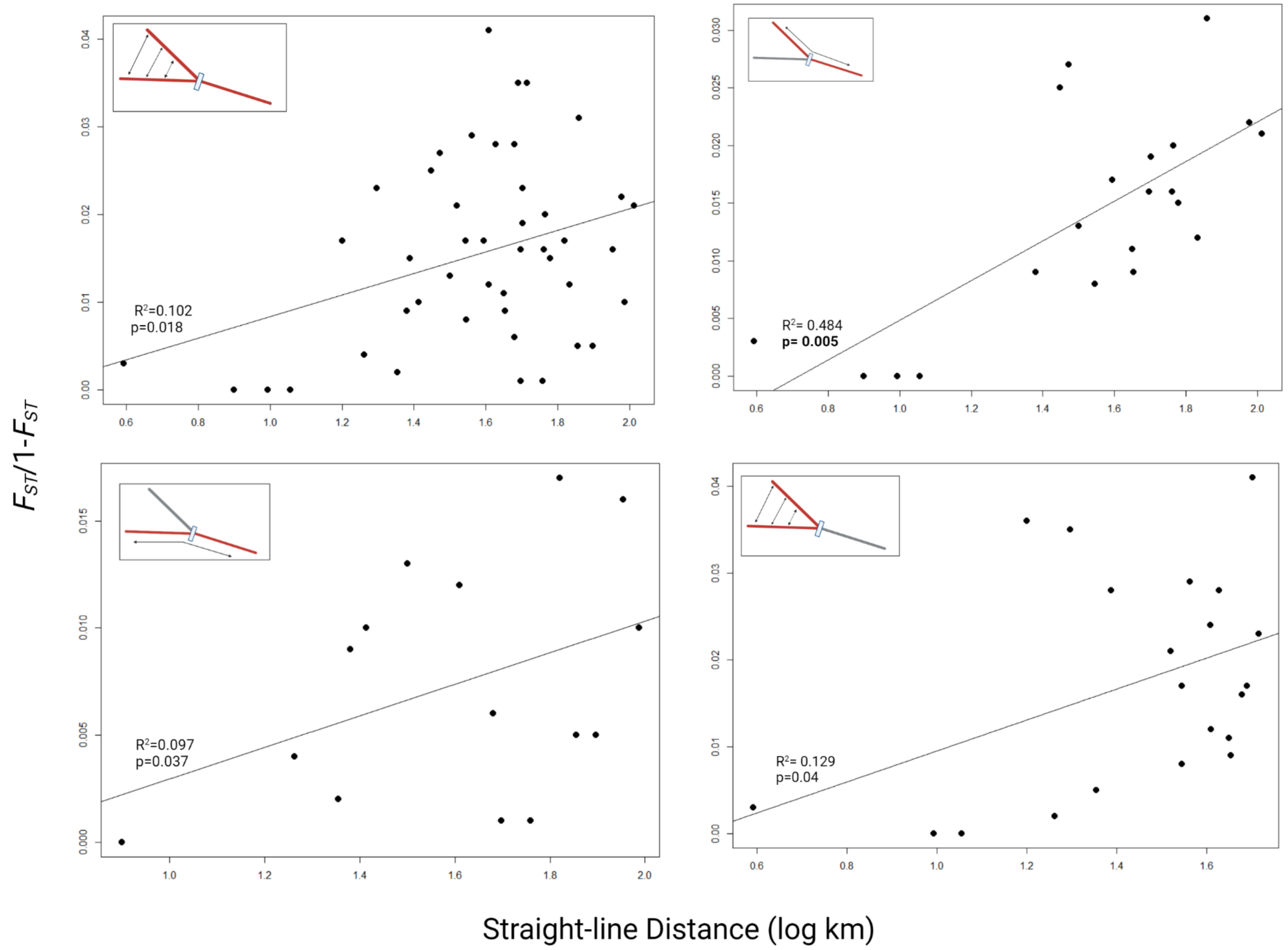

R²=0.102
p=0.018
R²= 0.484
p= 0.005
R²=0.097
p=0.037
R²= 0.129
p=0.04
FST/1-FST
Straight-line Distance (log km)

**Figure S2**. Mantel test for Isolation by distance along straight-line distance pathways. Straight line distance was log transformed.
